# Self-Organizing Map Methodology for Sorting Differential Expression Data of MMP-9 Inhibition

**DOI:** 10.1101/586628

**Authors:** Rachel St. Clair, Michael Teti, Ania Knapinska, Gregg Fields, William Hahn, Elan Barenholtz

## Abstract

An unsupervised machine-learning model, based on a self-organizing map (SOM), was employed to extract suggested target genes from DESeq2 differential expression analysis data. Such methodology was tested on matrixmetalloproteinase 9 (MMP-9) inhibitors. The model generated information on several novel gene hits that may be regulated by MMP-9, suggesting the self-organizing map method may serve as a useful analytic tool in degradomics research for further differential expression data analysis. Original data was generated from a previous study, which consisted of quantitative measures in changes of levels of gene expression from 32,000 genes in four different conditions of stimulated T-cells treated with an MMP-9 inhibitor. Since intracellular target of MMP-9 are not yet well characterized, the functional enrichment analysis program, WebGestalt, was used for validation of the SOM identified regulated genes. The proposed data analysis method indicated MMP-9’s prominent role in biological regulatory and metabolic processes as major categories of regulation of the predicted genes. Both fields suggest extensive intracellular targets for MMP-9-triggered regulation, which are new interests in MMP-9 research. The methodology presented here is useful for similar knowledge and discovery from quantitative datasets and a proposed extension of DESeq2 or similar data analysis.

## I. Introduction

**D**EGRADOMICS is the study of the cellular processes of proteases. Many analysis techniques employed in this field, such as DESeq2, generate statistical analyses of raw proteomics and genomics differential expression data, but fail to set a threshold clearly distinguishing regulated genes from non-regulated genes. Often, these approaches require laborious processing by domain experts and hand-selected parameters. Here, a novel unsupervised machine learning and data-processing technique, based on Self-Orgnaizing Maps (SOM) is introduced to automatically identify probable gene targets based on differential expression analysis data. To do so, the model finds points in the high-dimensional dataspace which can be interpreted as prototype representations of up-regulated, down-regulated, and non-regulated genes; distances from real datapoints (i.e. from actual genes) to these prototype points serve as a measure of the degree and type of gene regulation affected by treatment conditions.

In order to develop and test this model, differential gene expression data was collected from MMP-9 inhibitor treated T-cells. Recent interest in the family of matrix metalloproteinases (MMPs) has increased based on its pathogenic characteristics in many diseases. A major report reviews the role of a specific subset of MMPs, MMP-9, in multiple sclerosis, yet there is no clear consensus on the identity of the MMP-9 substrate pool [1]. In addition to direct proteolysis of substrates, the indirect role of MMP-9 on regulation of other targets still remains unknown. Intracellular targets for MMPs are not well defined, have not been studied extensively, and have only begun to be elucidated in recent years. Differential gene expression data generated by DESeq2 could potentially help to develop novel MMP-9 inhibitors.

In the current study, DESeq2 analysis of data collected from T-cells treated with MMP-9 inhibitor were used to train the SOM model in order identify highly regulated genes. Since intracellular substrates of MMP-9 are not well known, validation of the proposed model’s suggested regulated gene targets were given to a functional gene analysis program, WebGestalt. Results indicate that this method applied to differential expression analysis can result in effectively identifying target substrates and may serve as a more effective pipeline to investigate potential gene hits.

### A. Self-Organizing Maps

A self-organizing map was used as the major backbone of the model. The idea behind this class of models is to represent the inputs (e.g. genes and respective quantitative values) as nodes in coalesced sectors of the topographical data distribution [2], [3]. However, if the number of nodes is restricted to less than the count of inputs, the map is forced to organize the inputs into clustered bins. Since nodes model inputs, their associated tensor (data array) contains what can be described of as the ‘best-fit’ input for that particular node. Each tensor contains the best-represented values of the inputs’ features particular to each respective node. This unsupervised learning approach learns to model the inputs by multiplying initially random weights according to a loss function (distance of nodes from data). The objective of using a SOM in this application was to separate the genes of interest into more definable categories of expression.

### B. DESeq2

DESeq2 is an analysis program that ranks deferentially expressed genes from collected RNA-sequence data. The program is state-of-the-art for determining features of expression comparison from shrinkage estimators. Often, as in the current case, the resulting analysis consist of each sequenced gene corresponding to a number of features, most notably p-value and fold change [4]. Although a central aim of DESeq2 is to better visualize and prioritize a ranked gene list, manually sorting of genes based on different feature values is typically required which can be a highly time-intensive process. In this case, the task is not as easy as taking the lowest p-value and the highest fold change due to the possibility of confounding regular cellular processes with MMP-9 inhibitor effects (due to activated T-cell stimulation). Thus, the current approach proposes to extend data analysis of DESeq2 by automatically evaluating its output for different treatment condition of genes, based on their respective quantitative features.

### C. WebGestalt

WebGestalt is an online functional enrichment analysis program [5]. The major functional comparison algorithm within WebGestalt that was used in this study was the overrepresentation enrichment analysis which generates p-values calculated based on multiple-test correction enrichment of different functional categories. The resultant p-values generated by WebGestalt are used as a measure of confidence in a given set of genes functionality relationship. As used in this work, the genes selected by the model as likely to be regulated were given to Webgestalt for validation.

## II. Data

DEseq2 differential expression data was taken from prior experiments in which M. muscus stimulated T-cells were treated with MMP-9 inhibitor or vehicle (control). The resultant data yielded approximately 8,000 differentially expressed genes of interest. According to DESeq2 protocol, base mean read count, log2fold change, standard error of log2fold change, Wald statistic, p-value, and p-adjusted values were calculated for each of the following four conditions: treatment vs. vehicle at 3 hours, treatment vs. vehicle at 6 hours, treatment at 3 hours vs treatment at 6 hours, and vehicle at 3 hours vs vehicle at 6 hours. The resultant genes of interest dataset comprised 36,000 rows of 7 columns: gene name, base mean read count, log2fold change, standard error of log2fold change, Wald statistic, p-value, adjusted p-value, and condition. For the final input, we withheld gene name, based on preliminary testing which found this feature to be non-predictive.

## III. Model

Structure of the proposed model entails three distinct processes before a final report of predicted target genes is produced: gene regulation, gene result selection by differential algorithm, gene analysis using WebGestalt and literature verification. Most important is the adapted self-organizing map and differential comparisons (fig. 1) [6].

**Figure 1.**
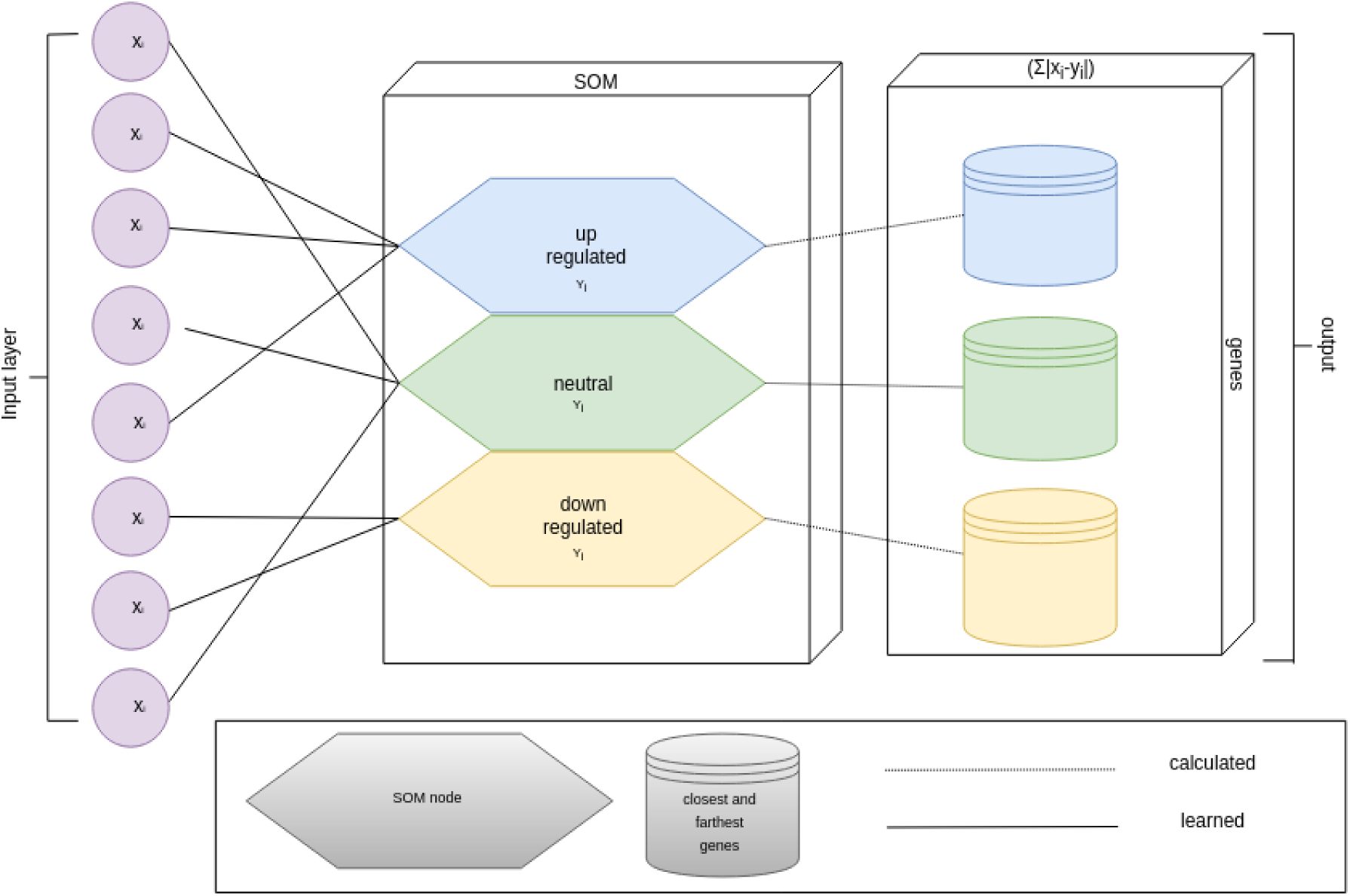
Basic network architecture for the self-organizing map and differential algorithm.

### 1) Gene Regulation Modeled by Nodes

For the SOM, three nodes were assigned to represent all 36,000 genes with the intent the map would organize the genes into upregulated, downregulated, and non-regulated categories. These labels were not assigned to the given inputs, but discerned after training. Identifiable by the log2fold change feature column, each node depicted the three intended categories, with positive, negative, or approximately zero numbers corresponding to upregulated, downregulated, and non-regulated, respectively. Through iterative training, weights of the associated nodes are updated according to a greedy policy, moving the nodes in a direction that best represents all the data-points. For each of the four conditions, the SOM learns to organize the condition’s genes into three nodes (eg. each condition has its own set of nodes). This technique effectively associated each of the genes of interest with one node, allowing for later comparisons.

### 2) Gene Result Selection by Differential Algorithm

To determine target genes, two comparisons needed to be made: 1 assess which genes of interest in each condition were being most regulated and 2 determine how genes of interest from a particular condition compared with an opposing condition’s nodes. To complete the first task, a measurement of how close each gene was to its respective condition’s nodes was needed. The second task required the same measurement from each gene of one condition to the nodes of a different condition. Thus, a distance algorithm was constructed to compare how similar a single data-point was to particular node (fig. 2). This function was applied to each possible within condition and between condition comparisons (fig. 3). In doing so, a list of most to least similar genes to nodes was created. The top most and least representative gene of each node (smallest and highest difference, respectively) per comparison was collected. Since the nodes are learned in a somewhat stochastic manner, variations in top regulated genes varied. To account for this variation, the entire above process was repeated for 1,000 epochs resulting in 1,000 genes of highest and 1,000 genes of lowest similarity for each node per compared conditions.

**Figure 2.**
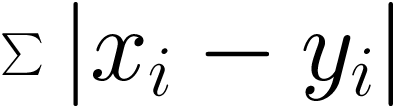
This function was used to assess the similarity between a particular node and data-point by subtracting a node vector (*y*_*i*_) from a data-point vector (*x*_*i*_).

**Figure 3.**
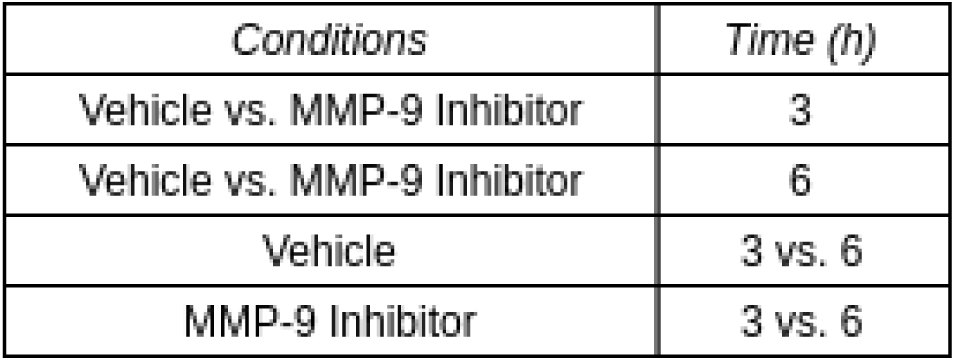
Comparisons of each node to each datapoint from these groups are essential in identifying MMP-9 possible targets.

### 3) Gene Validation Using WebGestalt

This master list is then sorted based on number of occurrences and reduced to the top 10 occurring genes in each category. Priority genes were collected and sorted from these categories by genes most occurring in comparisons between MMP-9 inhibitor vs. vehicle and MMP-9 inhibitor/vehicle at 3 vs. 6 hours, as these are the most straightforward comparisons of MMP-9 inhibitor effects (fig. 4). The list of 36 resultant genes were analyzed by Webgestalt enrichment program in Wikipathway (p=0.018) using parameters: mmusculus, overrepresentation enrichment analysis, pathway, wikipathway, genesymbol, genome (fig. 4). The full 36 suggested target gene list can be found in Appendix A.

**Figure 4.**
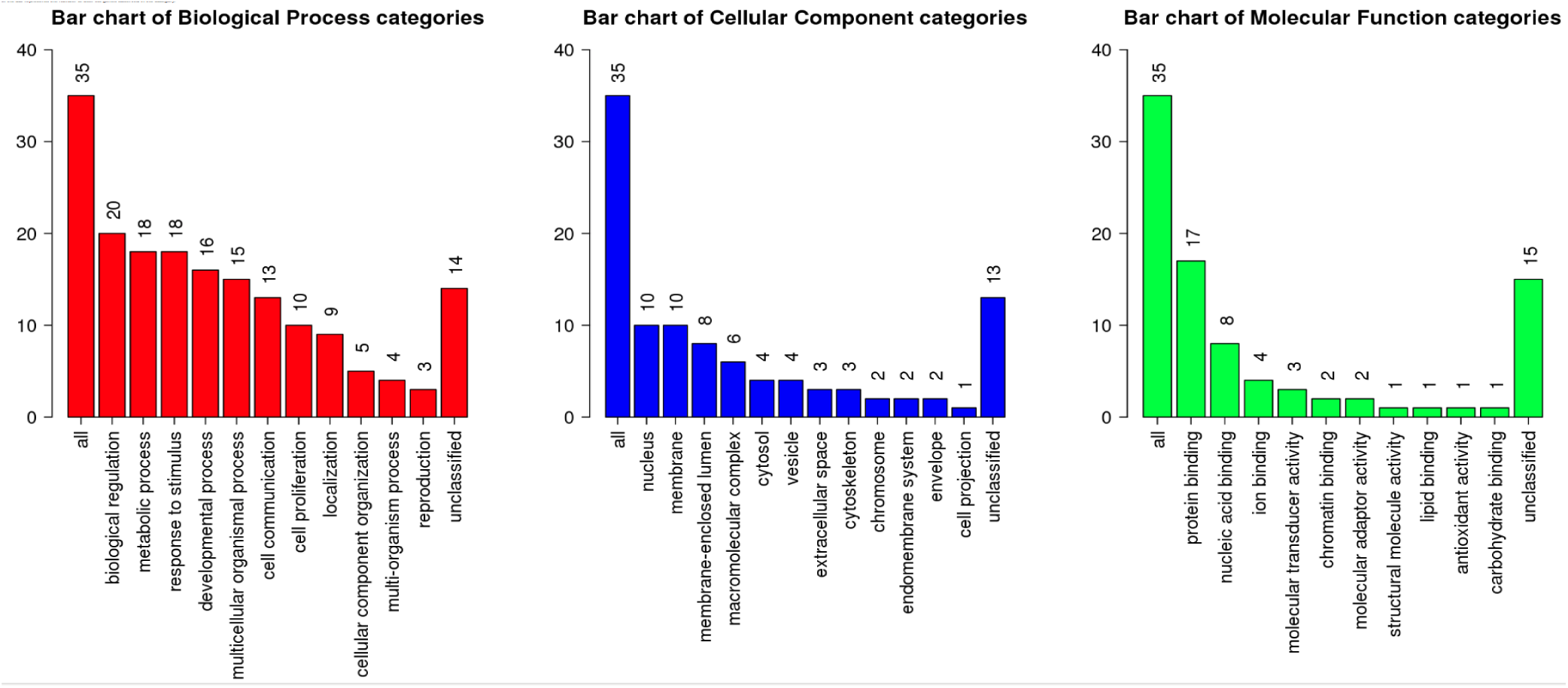
The resultant functional pathway associations for the 36 genes suggested as targets for MMP-9

## IV. Conclusion

This work will be utilized in efforts of an ongoing trial to produce effective MMP-9 inhibitors. Predicted targets will be validated as downstream effectors of MMP-9 intracellular function. Furthermore, the proposed model is lightweight and easily adaptable to different types of quantitative input (i.e. genomics or proteomics). Since the model requires almost no pre-processing if using DESeq2 data, it is readily applicable to such domains. Finally, these results suggest an even furthered automatic pipeline for data-analysis, decreasing the tedious and often confounding error propagated by human means. It can be foreseen that unsupervised models, such as SOM-MMP9, will gain more notoriety in related fields of inquiry.

### Appendix A

#### Suggested 36 Gene Target List

Malat1 Neat1 Gm26917 Gtf2ird1 Ifi27l2a Dll4 Elmsan1 Spata13 Fosl2 Ccnd2 Jag1 Nqo1 Il2 Lif Cd40 Trp53inp1 Gramd1b Serpinb9 Irf7 Nr4a1 Katna1 Dock10 Tkt Smek2 Junb Atp6v1d Rel Nfkbid Fscn1 Med14 Ccr7 Macf1 Spag9 Ccl22 Txnrd1 Egr1

### Appendix B

#### Code for SOM-MMP9

BLINDED FOR PEER-REVIEW https://github.com/MedBios/DeepProteins/tree/master/SOMMMP9

## Acknowledgment

We would like to thank the Machine Perception and Cognitive Robotics Laboratory for their use of Nvidia graphics processing units and Adam Ishay for his insightful help in debugging the model. This research was supported by a Neuroscience Pilot Award from the FAU Brain Institute to Ania Knapinska.

